# Lipid moieties of sonic hedgehog are important for interaction with its inhibitor, WIF1

**DOI:** 10.64898/2026.02.23.707386

**Authors:** Krisztina Kerekes, Mária Trexler, László Bányai, László Patthy

## Abstract

It has been recognized a long time ago that the hedgehog (Hh) and Wnt signaling pathways have numerous similarities that suggest their common evolutionary origin. Although the Hh and Wnt proteins are unrelated they are similar in as much as they carry lipid modifications that are critical for their interaction with their receptors. In our earlier work we have shown that Wnt inhibitory factor 1 (WIF1), originally identified as a Wnt antagonist also binds to and inhibits the signaling activity of sonic hedgehog (Shh), raising the possibility that the lipid moieties of these unrelated morphogens play a dominant role in their interaction with WIF1. In the present work we have compared the interactions of human WIF1 protein with lipidated and non-lipidated forms of human sonic hedgehog (Shh) using Surface Plasmon Resonance spectroscopy and reporter assays monitoring the signaling activity of human Shh. Our studies have shown that human WIF1 protein has significantly higher affinity for lipidated than non-lipidated Shh, indicating that lipid modifications of Hhs are important for interactions with WIF1.

## 1. Introduction

The hedgehog (Hh) and Wnt signaling pathways have numerous similarities that suggest their common evolutionary origin [1-3]. For example, signaling of the Hh and Wnt morphogens is mediated by closely related heptahelical receptors (Smoothened and Frizzled, respectively) and both pathways use GSK3β, CK1 and β-TrCP to regulate the proteolysis of the key transcriptional effectors of these pathways [1-3]. A surprising additional connection of the two pathways came with the discovery that Wnt Inhibitory Factor 1 (WIF1), originally identified as a Wnt antagonist, is also involved in the regulation of the activity of Hh [4-8]. In our earlier work we have shown that human WIF1 protein binds human sonic hedgehog (Shh) with high affinity and inhibits its signaling activity efficiently [9].

Although the protein components of Hhs and Wnts are unrelated, they are similar in that both morphogens are lipid-modified and that the lipid modifications are critical for their signaling activity.

Wnt proteins are palmitoleoylated in their N-terminal parts and this modification has been shown to be important for their signaling activity [10-11]. Analysis of the structure of Wnt8 in complex with the receptor, Frizzled-8, has provided an explanation for the importance of this lipid modification: the palmitoleic acid side-chain is inserted into a deep groove of the ligand-binding Fz domain of the receptor [12].

The precursors of Hh proteins are modified by palmitic acid at the very N-terminal end of the proteins after their signal sequence has been removed. The C-terminal protease domain of the Hh precursor cleaves the precursor in an autocatalytic manner to release the active N-terminal signaling domain and during this cleavage the C-terminus of the signaling domain becomes covalently modified by a cholesterol molecule [13-15]. Fatty-acylated Hh is far more active than the unacylated ligand, blocking Hh palmitoylation interferes with Hh signaling [16-18]. Recent studies on “native”, dually lipidated Hh in complex with its receptor, PTCH1, have shown that the morphogen grasps the extracellular domain of its receptor with two lipidic pincers, the N-terminal palmitate and the C-terminal cholesterol, that are both inserted into the PTCH1 protein core [19-20].

In view of the fact that the Hh and Wnt proteins are unrelated but both of them are lipid-modified it was plausible to assume that inhibition of the activities of these unrelated proteins by WIF1 might be mediated by interactions of WIF1 with the lipid moieties of these morphogens [3].

Recent studies have provided evidence for a critical role of lipid modifications of Wnts in their interaction with WIF1. de Almeida Magalhaes and coworkers have shown, that WIF1 (as well as secreted frizzled-related proteins) are Wnt carriers, these carriers form complexes with Wnts in a lipid-dependent manner [21]. It is worthy of note that in our earlier studies we have identified a hydrophobic site on the WIF-domain that was suggested to bind the lipid moiety of Wnts [22-23]. We have pointed out that the high affinity of WIF domains for palmitoleoylated groups would also explain why human WIF1 has remarkably high affinity for distantly related human Wnt paralogs, as well as Wnts from a variety of mammals, *Xenopus* and *Drosophila* [22].

As a working hypothesis we have assumed that the lipid moieties of Hhs may also bind to the hydrophobic binding site of the WIF domain and these interactions may also be critical for the inhibitory activity of WIF1 on Hh signaling [3]. To test this assumption, in the present work we have compared the interactions of human WIF1 protein with lipidated and non-lipidated forms of human sonic hedgehog (Shh) using Surface Plasmon Resonance spectroscopy, pull-down experiments and reporter assays monitoring the signaling activity of human Shh. These studies have shown that both lipidated and non-lipidated Shh had pronounced affinity for human WIF1 protein, but the lipidated form had significantly higher affinity, indicating that lipid modifications are important for interactions with WIF1.

## 2. Materials and Methods

### 2.1. Proteins, cell lines, media and reagents

Recombinant human WIF1 (rhWIF1), recombinant dually lipidated human sonic hedgehog protein expressed in HEK cells (rhShh_8908-SH) and non-lipidated human sonic hedgehog proteins expressed in *E. coli* (rhShh_1314-SH and rhShh_1845-SH) were purchased from R&D Systems (Minneapolis, MN USA). The protein sequences of rhShh_8908-SH and rhShh_1314-SH correspond to residues Cys24-Gly197 of human sonic hedgehog, whereas rhShh_1845-SH differs from these proteins in that the Cys24 residue has been replaced by a hydrophobic Met-Ile-Ile sequence to mimic the hydrophobic palmitoyl moiety of dually lipidated sonic hedgehog protein.

Human WIF1 Antibody (Goat IgG, AF134) and Human/Mouse Sonic Hedgehog/Shh N-Terminus Antibody (Goat IgG, AF464) were from R&D Systems (Minneapolis, MN USA). Anti-Goat IgG (whole molecule)-Alkaline Phosphatase antibody produced in rabbit (A4187) and Anti-Rat IgG (whole molecule)–Alkaline Phosphatase antibody produced in goat were from Sigma-Aldrich (St Louis, Mo USA). The ONE-Step™ Luciferase Assay System and the Gli Reporter-NIH3T3 cell line were purchased from BPS Bioscience (Inc., San Diego, CA, USA). Opti-MEM Reduced Serum Medium (Gibco by Thermo Fisher Scientific, Waltham, MA, USA) with 0.5% calf serum (Merck - SigmaAldrich, KGaA, Darmstadt, Germany), 1% non-essential amino acids (Gibco by Thermo Fisher Scientific, Waltham, MA, USA) and 1% penicillin/streptomycin was used for reporter assays. Culture media DMEM was obtained from Merck - SigmaAldrich (KGaA, Darmstadt, Germany). Luminescence was measured using an EnSpire plate reader (PerkinElmer, Inc. Waltham, MA, USA).

Pierce™ Classic Immunoprecipitation Kit (26146, Thermo Fisher Scientific, Waltham, MA USA) was used for pull-down assays.

CM5 sensor chips and the reagents for protein coupling to the chips were from Biacore AB (Uppsala, Sweden). Amersham Protran Premium nitrocellulose blotting membrane was from GE Healthcare Life Sciences (Marlborough, Massachusetts, USA). Nitro Blue tetrazolium and 5-bromo-4-chloroindol-2-yl phosphate were from Serva Electrophoresis (Heidelberg, Germany).

### 2.2. Pull-down assays

The rhWIF1 protein (12 pmole) was incubated with rhShh_1314-SH or rhShh_1845-SH protein at a 1:1 molar ratio for 1 h at 4 °C in TBS buffer (25 mM Tris, 150 mM NaCl; pH 7.2; final volume 30 μL). Antibodies against rhWIF1 or rhShh were then added to the respective mixtures at a molar ratio of 1:1.5, and the solutions (final volume 50 μL) were further incubated for 1 h at 4 °C. The resulting protein–protein–antibody complexes (50 μL) were applied to microcentrifuge columns containing 45 μL Pierce Protein A/G Plus Agarose. After adding 100 μL TBS buffer, the columns were gently agitated overnight at 4 °C. Subsequently, the columns were centrifuged, the A/G Plus Agarose was washed four times with 200 μL TBS buffer, and the bound proteins were eluted with 50 μL elution buffer (pH 2.8), following the manufacturer’s protocol. The eluates were analyzed by Western blotting using antibodies specific for WIF1 or Shh proteins.

### 2.3. Surface plasmon resonance analyses

SPR measurements were performed on a BIACORE X (GE Healthcare, Stockholm, Sweden) instrument. Proteins to be immobilized were dissolved in 50 mM sodium acetate buffer, pH 4.5, and solutions were injected with a 5 μL/min flow rate on a CM5 sensor chip (Cytiva, Uppsala, Sweden) activated by the amine coupling method, according to the manufacturer’s instructions. For interaction measurements, 90 μL aliquots of protein solutions were injected over the sensor chips with a 20 μL/min flow rate. Binding and washes were performed in 20 mM Tris, 150 mM NaCl, 5 mM EDTA, 0.005% Tween 20, 100 μM CHAPS, pH 7.4 buffer. After each cycle, the chips were regenerated by injection of 35 µL of 8 M urea, 1M NaCl, 100 mM Tris, 5 mM EDTA, and 0.005% Tween-20, pH 7.4. Control flow cells were prepared by performing the coupling reaction in the presence of coupling buffer alone. Control flow cells were used to obtain control sensorgrams showing nonspecific binding to the surface as well as refractive index changes resulting from changes in bulk properties of the solution. Control sensorgrams were subtracted from sensorgrams obtained with immobilized ligand.

All experiments were repeated at least three times. To correct for differences between the reaction and reference surfaces, we also subtracted the average of sensorgrams obtained with blank running buffer injections. The kinetic parameters for each interaction were determined by fitting the experimental data with BIAevaluation software 4.1 and the closeness of the fits was characterized by the χ2 values. Only fits with χ2 values lower than 5% of the Rmax were accepted. Data were fitted to a model of 1:1 Langmuir interaction.

### 2.4. Reporter assays

The signaling activities of human sonic hedgehogs and the hedgehog antagonist activities of WIF1 protein were evaluated using the Gli Reporter-NIH3T3 cell line, which carries the firefly luciferase gene under the control of Gli-responsive elements stably integrated into NIH3T3 cells.

For reporter assays monitoring the signaling activities of sonic hedgehogs, 2.5 × 10^4^ cells/100 μL/well were seeded in 96-well tissue culture plates and incubated for ∼24 h in DMEM/glutamine/calf serum/penicillin/streptomycin medium at 37 °C in a CO_2_ incubator. When cells reached confluency, the culture medium was removed, and 50 μL aliquots of serial dilutions of rhShh_8908-SH (0–600 nM), rhShh_1314-SH (0–2400 nM) or rhShh_1845-SH (0–2400 nM) prepared in assay medium (Opti-MEM Reduced Serum Medium supplemented with 0.5% calf serum, 1% non-essential amino acids, and 1% penicillin/streptomycin) were added to the wells. Plates were incubated for 18 h, after which luminescence was measured using an EnSpire plate reader.

For reporter assays measuring the hedgehog antagonist activities of WIF1 protein, cells were grown to confluency, the medium was removed, and 50 μL aliquots containing 15 nM rhShh_8908-SH or rhShh_1314-SH or rhShh_1845-SH, preincubated for 5 min with 0–1000 nM rhWIF1 in assay medium, were added to the wells.

For control experiments, confluent cells were treated with 50 μL aliquots of assay medium only (unstimulated control wells). To determine background luminescence, 50 μL aliquots of assay medium were added to cell-free control wells.

Each experiment was performed three times, with three replicates per condition.Luciferase assays were carried out using the ONE-Step™ Luciferase Assay System according to the manufacturer’s instructions: 50 μL of ONE-Step™ Luciferase reagent was added to each well, and the plates were shaken for 20 min at room temperature. Luminescence was measured using EnSpire plate reader. Background luminescence (from cell-free control wells) was subtracted from all measurements, and the average fold induction of Gli luciferase reporter expression was calculated by comparing the luminescence of stimulated and unstimulated wells.

Dose–response data for the signaling activities of human sonic hedgehogs proteins and WIF1-mediated inhibition of the signaling activities of sonic hedgehogs proteins were analyzed using a four-parameter Hill (logistic) model implemented in the Nonlinear Curve Fit module of OriginPro 2025.

Normalized luminescence values were plotted against the logarithm of ligand concentration, and parameters were optimized via nonlinear least-squares fitting using the Levenberg–Marquardt algorithm, yielding EC_50_ values for increase or decrease of signaling activity. The Hill coefficient (nH) was extracted from the fitted curves to characterize the steepness and apparent cooperativity of the response.

Pairwise comparisons of the fitted dose-response curves of the activity (or inhibition) of the different recombinant sonic hedgehog proteins were performed using an ANOVA-based Extra sum-of-squares F-test of OriginPro 2025.

### 2.5. Protein analyses

For Western blotting, samples were separated on a non-reducing 16% SDS gel and subsequently transferred onto nitrocellulose membranes. Membranes were blocked for 1 h at room temperature in 10 mM Tris/HCl, 150 mM NaCl, and 0.05% Tween-20, pH 7.5 (TBST), supplemented with 5% non-fat dry milk. Primary antibody (0.2 μg/10 mL) diluted in TBST was applied for 2 h at room temperature, followed by three washes with TBST. Secondary antibodies, diluted 30,000-fold in TBST, were then incubated with the membranes for 1 h at room temperature, after which the blots were washed again three times with TBST. Protein detection was performed by immersing the membranes in 100 mM Tris/HCl, 100 mM NaCl, 5 mM MgCl_2_, 0.5 mM Nitro Blue tetrazolium, and 0.5 mM 5-bromo-4-chloroindol-2-yl phosphate (pH 9.5).

### 2.6. Statistical analyses

The statistical package of Origin 2018 was used for all data processing and statistical analysis. Statistical significance was set as a p value of < 0.05.

## 3. Results and Discussion

### 3.1. Human Wnt Inhibitory Factor 1 binds non-lipidated human Shh proteins with high affinity

Pull-down experiments illustrated in **Figure 1** have revealed that human WIF1 forms stable complexes with human Shh expressed in *E. coli*, despite the absence of lipid moieties.

**Figure 1.**
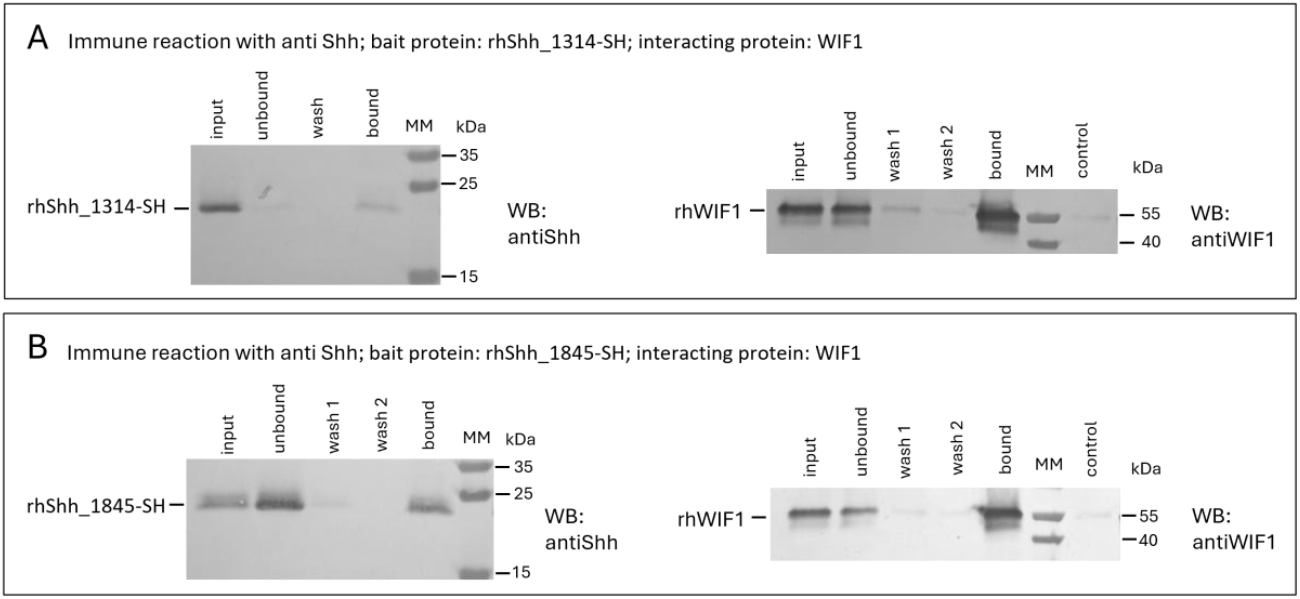
Human WIF1 binds non-lipidated human Shh proteins. In pull-down assays, rhWIF1 was incubated with *E. coli*-expressed rhShh_1314-SH (**Panel A**) or rhShh_1845-SH (**Panel B**), mixed with anti-Shh, and applied to Pierce Spin Columns containing Pierce Protein A/G Plus Agarose. Bound proteins were eluted with Pierce Classic IP kit Elution Buffer (pH 2.8), analysed by SDS/PAGE, and detected by Western blotting (WB) with antibodies against rhWIF1 and rhShh. As control, rhWIF1 was incubated with anti-Shh and analysed similarly. Lanes: input, sample applied to the column; unbound fraction; washing fractions (wash1, wash2); bound fraction; MM, PageRuler Plus Protein Ladder; control, as described above. Numbers indicate the molecular mass of the protein ladder bands.

In order to obtain quantitative information about the affinity of rhShh_1314-SH for WIF1, we have used Surface Plasmon Resonance spectroscopy measurements. When we used immobilized human Wnt Inhibitory Factor 1 as ligand, injection of human rhShh_1314-SH solutions elicited SPR signals in a dose-dependent manner; representative experiments are shown in **Figure2**. Analyses of the SPR response curves of these experiments have revealed that the K_d_ value of the WIF1-rhShh_1314-SH complex is 5.44 ± 0.41 nM (**Table 1**).

**Table 1.** Comparison of the kinetic parameters of the interactions of rhShh_1314-SH and human rhShh_8908-SH with immobilized human WIF1 protein (WIF1). The rate constants of the association and dissociation reactions and the equilibrium dissociation constants of the interactions were determined from surface plasmon resonance measurements with the BIAevaluation software 4.1.

| Interacting proteins | Kd (nM) | ka ($10^4 \times 1/\text{Ms}$) | kd ($10^{-4} \times 1/\text{s}$) |
| --- | --- | --- | --- |
| rhShh_1314-SH – WIF1* | $5.44 \pm 0.41$ | $3.86 \pm 0.24$ | $2.1 \pm 0.09$ |
| rhShh_8908-SH – WIF1** | $2.06 \pm 0.75$ | $6.85 \pm 0.6$ | $1.41 \pm 0.5$ |

\* Present paper
\*\* Published in Kerekes et al., Cells, 2021, 10(12):3496 [9]

**Figure 2.**
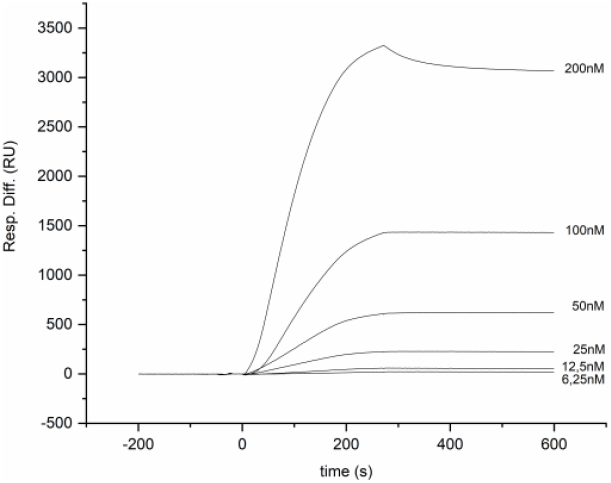
Characterization of the interaction of human rhShh_1314-SH with human WIF1 by surface plasmon resonance spectroscopy. Various concentrations of rhShh_1314-SH (6.25 nM, 12.5 nM, 25 nM, 50 nM, 100 nM and 200 nM) were injected over CM5 sensor chips containing immobilized rhWIF1 (at 0s on the abscissa). Note that the SPR response increases parallel to the increase in Shh concentration.

It should be pointed out that although these experiments suggest high affinity of WIF1 for rhShh_1314-SH (K_d_ =5.40 ± 0.40 nM), this affinity is significantly (p < 0.05) lower than for dually lipidated rhShh_8908-SH (K_d_ =2.06 ± 0.75 nM, see **Table 1**).

### 3.2. Human Wnt Inhibitory Factor 1 is a less potent antagonist of the signaling activity of non-lipidated than dually lipidated human Shh protein

#### 3.2.1. Signaling activities of non-lipidated and dually lipidated human Shh protein

In order to assess the role of lipid modifications of Shh in its interaction with human WIF1 we have used a reporter assay to monitor the effect of WIF1 on the signaling activity of rhShh_8908-SH, rhShh_1314-SH and rhShh_1845-SH. The Gli Reporter-NIH3T3 cell line used in this assay contains the firefly luciferase gene under the control of Gli responsive elements stably integrated into NIH3T3 cells. Activation of the hedgehog pathway by binding of Shh to the Patched receptor correlates with luciferase expression.

In this reporter assay the signaling activities of non-lipidated human Shhs expressed in *E. coli* were only moderately inferior to that of dually lipidated Shh, expressed in HEK cells (**Figure 3, Table 2**). Recombinant protein, rhShh_8908-SH, rhShh, corresponding to residues Cys24-Gly197 of human sonic hedgehog, cholesterol-modified at the C-terminal and fatty acid-modified at the N-terminal, increased luminescence ∼80-fold with an EC_50_ value of 34.22 ± 2.25 nM (**Figure 3A**), whereas the non-lipidated form of the same protein sequence, rhShh_1314-SH increased expression of the luciferase ∼ 70-fold with an EC50 value of 64.81 ± 1.29 nM (**Figure 3 B**). The non-lipidated recombinant protein rhShh_1845-SH that differs from rhShh_1314-SH in that the Cys24 is replaced by a hydrophobic Met-Ile-Ile sequence (intended to mimic palmitoylation of Shh), increased luminescence of reporter cells ∼60-fold with an EC50 value of 86.09 ± 1.11 nM (**Figure 3 C**).

**Table 2.** Parameters of the fit of the four-parameter Hill-equation to sigmoidal dose–response curves for rhShh_8908-SH, rhShh_1314-SH, and rhShh_1845-SH. EC_50_ (nM): concentration producing half-maximal activation; 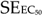: standard error of the EC_50_ estimate; nH: Hill coefficient describing curve steepness; SE_nH_: standard error of the Hill coefficient; R^2^ (COD): coefficient of determination indicating goodness of fit; N (repl.): number of replicates.

|  | EC<sub>50</sub> (nM) | SE<sub>EC<sub>50</sub></sub> | n<sub>H</sub> | SE<sub>n<sub>H</sub></sub> | R<sup>2</sup> (COD) | N (repl.) |
| --- | --- | --- | --- | --- | --- | --- |
| rhShh_8908-SH | 34.22 | 2.25 | 2.83 | 0.42 | 0.98 | 3 |
| rhShh_1314-SH | 64.81 | 1.29 | 1.59 | 0.18 | 0.99 | 3 |
| rhShh_1845-SH | 86.09 | 1.18 | 1.49 | 0.23 | 0.99 | 3 |

**Figure 3.**
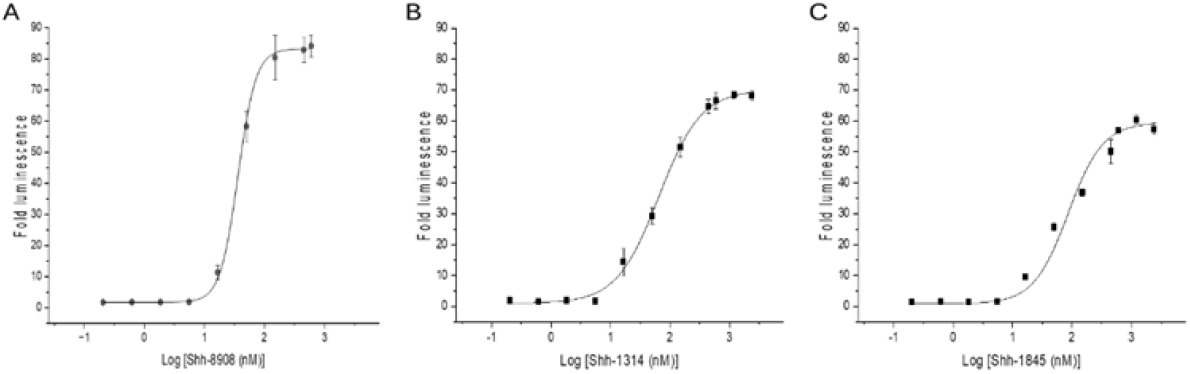
Comparison of the signaling activities of rhShh_8908-SH, rhShh_1314-SH and rhShh_1845-SH in the Gli Reporter-NIH3T3 cell line reporter assay. Panel A: rhShh_8908-SH expressed in HEK293 cells was applied at 0–600 nM. EC_50_ = 34.22 ± 2.25 nM. Panel B: rhShh_1314-SH expressed in *E. coli* was applied at 0–2400 nM. EC_50_ = 64.81 ± 1.29 nM. Panel C: rhShh_1845-SH expressed in *E. coli* was applied at 0–2400 nM. EC_50_ = 86.09 ± 1.11 nM. In all assays, Gli Reporter-NIH3T3 cells were grown to confluency, the culture medium was removed, and cells were treated with 50 μL aliquots of the indicated Shh concentrations. Data represent one experiment performed in triplicate and are expressed as fold induction relative to luminescence at 0 nM Shh.

Pairwise comparisons of the fitted dose-response curves of the activities of the different recombinant sonic hedgehog proteins revealed that the Extra sum-of-squares F-test detects statistically significant differences in dose–response profiles of the dually lipidated and non-lipidated morphogens (**Table 3**).

**Table 3.** Statistical significance of the differences between the dose–response curves of the three recombinant sonic hedgehog proteins, determined by the extra sum-of-squares F-test of pairwise comparisons.

| Sonic hedgehog proteins compared | p value |
| --- | --- |
| rhShh_8908-SH vs. rhShh_1314-SH | 0.000507 |
| rhShh_8908-SH vs. rhShh_1845-SH | 0.00625 |
| rhShh_1314-SH vs rhShh_1845-SH | 0.07339 |

It must be pointed out that the moderate (less than 3-fold) difference in the EC_50_ values of lipidated and non-lipidated Shh proteins in the reporter assay based on the Gli Reporter-NIH3T3 cell line is in striking contrast with the dramatic differences in their bioactivities measured by their ability to induce alkaline phosphatase production by C3H10T1/2 mouse embryonic fibroblast cells. As indicated by the manufacturer of the rhShh_8908-SH, rhShh_1314-SH and rhShh_1845-SH proteins, in the latter assay dually lipidated rhShh_8908-SH is over 250-fold more active than non-lipidated Shh with the same amino acid sequence and 14-fold more active than non-lipidated rhShh_1845-SH with a modified, more hydrophobic N-terminus. It is also noteworthy that although in the C3H10T1/2 mouse embryonic fibroblast cell assay rhShh_1845-SH (with a more hydrophobic N-terminus), is significantly more active than rhShh_1314-SH, in the reporter assay based on the Gli Reporter-NIH3T3 cell line, the difference was just the opposite (see **Figure 3, Table 2)**. The EC_50_ value of rhShh_1314-SH (64.81 ± 1.29 nM) was slightly but significantly lower (p<0.05) than that of rhShh_1845-SH (86.09 nM ± 1.11 nM).

The explanation for this apparent discrepancy may lie in the marked differences of the two types of assays. The C3H10T1/2 cell line is a murine pluripotent fibroblastic cell line that differentiates into an osteoblastic phenotype in the presence of Shh, a process that may be monitored by measuring alkaline phosphatase induction in the cultures following treatment with Shh [24, 25]. An important aspect of this assay is that the expression of the patched gene, the receptor of Shh, is upregulated several-fold by Shh in C3H10T1/2 cells [24, 26] and this may result in significant amplification of Shh signaling. The possible significance of this amplification may be illustrated by the observation that whereas modifications at the N-terminus of Shh dramatically increased or decreased the potency of Shh in the C3H10T1/2 cell line assay, the effects of these modifications were barely detectable in their binding affinity for the receptor protein, patched [26, 27]. Apparently, the Gli Reporter-NIH3T3 cell line based assay is suitable for monitoring Shh signaling through patched receptor, but lacks the signal amplification feature characteristic of the C3H10T1/2 cell line assay.

Comparison of the dose-response curves of rhShh_8908-SH, rhShh_1314-SH, and rhShh_1845-SH (**Figure 3, Table 2**), revealed that the fitted curve is significantly steeper and the Hill coefficient is significantly higher (**n**_**H**_ =2.83) in the case of dually lipidated Shh than non-lipidated Shhs, 1314-SH, (**n**_**H**_ = 1.59) and rhShh_1845-SH (**n**_**H**_ =1.49).

These observations indicate marked differences in the response of the patched receptor to lipidated and non-lipidated sonic hedgehog. The lipidated morhogen has both higher affinity (lower EC50 value) and increased sensitivity (higher nH value) than non-lipidated forms. The structural basis for the higher sensitivity of patched to lipidated than non-lipidated soninc hedgehog is not fully understood at present.

In a recent study Petrov and coworkers [27] have also found that dose–response curves of signaling by lipidated sonic hedgehog through patched receptor have steep slopes with Hill coefficient significantly greater than 1 (n ≈ 2), indicating that activation of hedgehog signaling by sonic hedgehog is switch-like. In principle, the switch-like response of wild-type patched to sonic hedgehog and a Hill coefficient of nH ≈ 2 could be explained by either allostery or ultrasensitivity. Since sonic hedgehog binds two patched molecules [28], coupling of receptor protomers via allostery could account for nH≈ 2. Petrov and coworkers [27], however, have pointed out that allosteric coupling cannot explain the nH ≈ 3 observed for some patched mutants. Furthermore, the authors have observed an nH ≈ 2 for the response to palmitoylated SHH22 peptide that can not dimerize patched as it lacks the globular domain of sonic hedgehog, leading the authors to suggest that ultrasensitivity is a likelier explanation for the switch-like activation of the Hh pathway than allostery.

Taken together, the fact that palmitoylated forms and fragments of Shhs have Hill coefficients significantly higher than non-lipidated Shhs suggests that palmitoylation of Shh has a key role in the switch-like activation of the Hh pathway by this morphogen.

#### 3.2.2. Signaling activity of dually lipidated Shh protein is more sensitive to inhibition by Wnt Inhibitory Factor 1 than the activities of non-lipidated Shh proteins

Our studies on the influence of human WIF1 on the signaling activities of Shhs have revealed significant (p<0.05) differences in the potency of the inhibitor (**Figure 4, Table 4**. Analyses of the dose-response curves of the inhibitor assays have shown that human WIF1 protein inhibits the signaling activity of the non-lipidated rhShh_1314-SH and rhShh_1845-SH proteins with EC50 values of 6.83 ± 0.89 nM and 15.71 ± 1.75 nM, respectively, whereas the EC50 value for dually lipidated rhShh_8908-SH was significantly lower, 3.78 ± 0.13 nM. These observations suggest that although lipidation of Shh is not indispensable for its interaction with WIF1 protein it contributes significantly to the stability of the WIF1-Shh complex.

**Table 4.** Parameters of the fit of the four-parameter Hill-equation to sigmoidal dose–response curves for the WIF1 inhibition of rhShh_8908-SH, rhShh_1314-SH, and rhShh_1845-SH. EC_50_ (nM): concentration producing half-maximal inhibition; 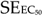: standard error of the EC_50_ estimate; nH: Hill coefficient describing curve steepness; SE_nH_: standard error of the Hill coefficient; R^2^ (COD): coefficient of determination indicating goodness of fit; N (repl.): number of replicates.

| | $EC_{50}$ (nM) | $SE_{EC_{50}}$ | $n_H$ | $SE_{n_H}$ | $R^2$ (COD) | N (repl.) |
| --- | --- | --- | --- | --- | --- | --- |
| rhShh_8908-SH + WIF1 | 3.78 | 0.13 | -2.60 | 0.31 | 0.99 | 3 |
| rhShh_1314-SH + WIF1 | 6.83 | 0.89 | -2.10 | 0.29 | 0.98 | 3 |
| rhShh_1845-SH + WIF1 | 15.71 | 1.75 | -1.95 | 0.21 | 0.99 | 3 |

**Figure 4.**
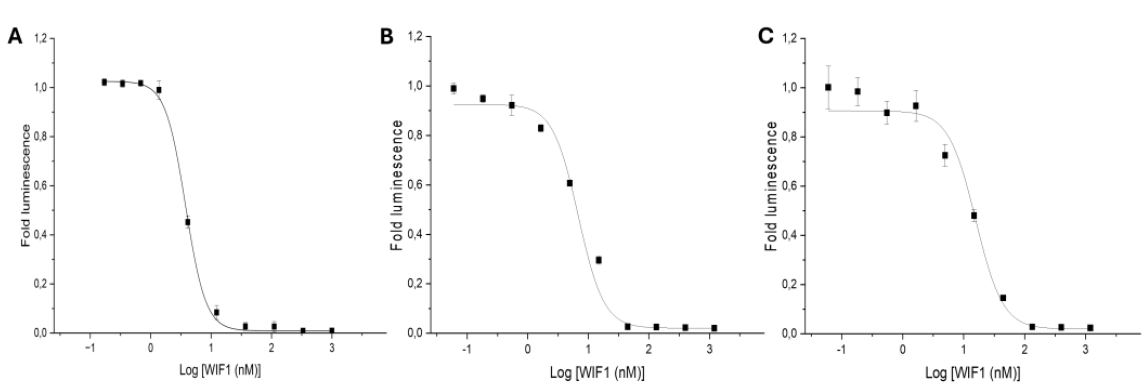
Inhibition of the signaling activities of rhShh_8908-SH, rhShh_1314-SH and rhShh_1845-SH proteins by human WIF1 protein in the Gli Reporter-NIH3T3 cell line reporter assay. Panel A: Inhibition of human rhShh_8908-SH expressed in HEK293 cells (15 nM) by rhWIF1 (0–1000 nM); EC_50_ = 3.78 ± 0.13 nM. Panel B: Inhibition of rhShh_1314-SH expressed in E. coli (15 nM) by rhWIF1 (0–1200 nM); EC_50_ = 6.83 ± 0.89 nM. Panel C: Inhibition of rhShh_1845-SH expressed in E. coli (15 nM) by rhWIF1 (0–1200 nM); EC_50_ = 15.71 ± 1.75 nM. In all assays, Gli Reporter-NIH3T3 cells were grown to confluency, culture medium was removed, and cells were treated with 50 μL aliquots of rhShh preincubated for 5 min with rhWIF1. Data represent representative experiments performed in triplicate (B, C) or quadruplicate (A) and are expressed as fraction of luminescence relative to 0 nM rhWIF1.

Analysis of the dose–response curves of the WIF1-mediated inhibition of Shh proteins revealed that these curves are steep, with Hill-coefficients ≥ 2, suggesting that inhibition is switch-like (**Figure 4, Table 4**). The fitted curve, however, is steeper and the Hill coefficient is higher (**n**_**H**_ =-2.60) in the case of dually lipidated Shh than in the case of non-lipidated Shhs, 1314-SH, (**n**_**H**_ = -2.10) and rhShh_1845-SH (**n**_**H**_ =-1.95), suggesting that lipid moieties play a role in the switch-like inhibition of Shh by WIF1. Pairwise comparisons of the fitted dose-response curves of the inhibition of the different recombinant sonic hedgehog proteins by WIF1 have confirmed that the Extra sum-of-squares F-test detects statistically significant differences in dose–response profiles of the dually lipidated and non-lipidated morphogens (**Table 5**).

**Table 5.**
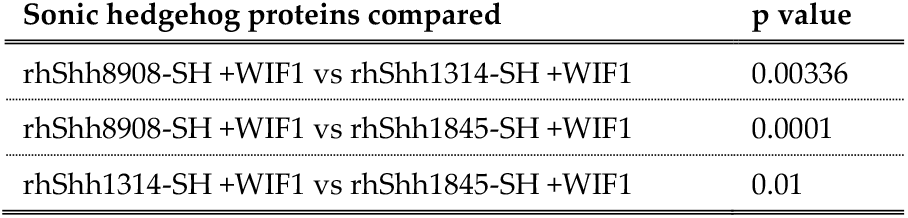
Statistical significance of the differences between the dose–response curves of the inhibition of recombinant sonic hedgehog proteins by WIF1, determined by the extra sum-of-squares F-test of pairwise comparisons.

| Sonic hedgehog proteins compared | p value |
| --- | --- |
| rhShh8908-SH +WIF1 vs rhShh1314-SH +WIF1 | 0.00336 |
| rhShh8908-SH +WIF1 vs rhShh1845-SH +WIF1 | 0.0001 |
| rhShh1314-SH +WIF1 vs rhShh1845-SH +WIF1 | 0.01 |

In summary, our observation that lipidated Shh is significantly more sensitive to inhibition by WIF1 than non-lipidated Shhs suggests that lipid moieties of this morphogen are important for interaction with its inhibitor, WIF1. It is also worth pointing out that WIF1 has significantly (p<0.05) lower affinity (higher EC50 value) for rhShh_1845-SH than for rhShh_1314-SH (**Table 4**). Since the two proteins differ only in that the N-terminal Cys24 residue of rhShh_1314-SH has been replaced by a Met-Ile-Ile sequence in rhShh_1845-SH, this difference is in harmony with the notion that the N-terminal arm of Shh (and its palmitoyl group) is involved in its interactions with WIF1.

## 4. Conclusions

We have shown earlier that Wnt inhibitory factor 1 (WIF1) protein binds human sonic hedgehog (Shh) with high affinity and inhibits its signaling activity [9]. Since hedgehog and Wnt proteins are unrelated but both of them are lipid-modified we have hypothesized that inhibition of the activities of these proteins by WIF1 might be mediated by interactions of WIF1 with the lipid moieties of the morphogens [3].

Our earlier studies on the interaction of WIF1 with Wnts have revealed that the N-terminal, lipid-modified part of Wnts interacts with an alkyl-binding site of the WIF domain of WIF1 [22, 23], therefore we have assumed that lipid moieties of Shh might also be involved in its interaction with WIF1 [3]. Furthermore, since the palmitoyl group of Shh is attached to the long, extended N-terminal arm of Shh we have assumed that this N-terminal, lipid-modified region may be critical for the Shh-WIF1 interaction [3].

To test these assumptions, in the present work we have compared the interactions of human WIF1 protein with lipidated and non-lipidated forms of wild type human Shh, as well as a variant of human Shh with a modified N-terminal sequence.

These studies have shown that both lipidated and non-lipidated wild type human Shh proteins have high affinity for WIF1, indicating that protein-protein interactions play a dominant role in the inhibition of Shh by the WIF domain. However, non-lipidated human Shhs were found to be significantly less sensitive to inhibition by WIF1, indicating that lipid modification plays an important role in the interaction of WIF1 with Shh.

Our finding that a variant of human non-lipidated Shh with a modified N-terminal sequence has significantly lower affinity for WIF1 than the wild type non-lipidated protein is consistent with the assumption that the N-terminal arm of human Shh is involved in its interaction with WIF1.

The main conclusion of the present work, that WIF1 is a more potent inhibitor of lipidated than non-lipidated sonic hedgehog, has important implications for the therapeutic application of this morphogen.

Recent studies have shown that there are several conditions where sonic hedgehog gene therapy or delivery of exogenous hedgehog protein has significant therapeutic potential. These include stimulation of peripheral nerve regeneration by Shh to treat spinal cord injury or stimulation of hair growth by Shh to treat hair loss [29-30]. It has been shown that sonic hedgehog may be used to protect brain or heart tissues from ischemic injury [31-32]. Significantly, delivery of exogenous hedgehog proteins has therapeutic potential for diabetic neuropathy and to treat impaired wound healing associated with diabetes [33-35]

We suggest that our finding, that lipidated Shh is more sensitive to inhibition by WIF1 than non-lipidated forms, must be taken into account in the design of Shh-based therapies of these diseases.

## Author Contributions

Conceptualization, L.P.; methodology, K.K., T.M. and L.B; software, L.B.; validation, K.K., L.B., T.M. and L.P.; formal analysis, K.K., L.B., T.M. and L.P.; investigation, L.P.; resources, L.P.; data curation, K.K., T.M. and L.B.; writing—original draft preparation, L.P.; writing—review and editing, K.K., L.B., T.M. and L.P.; visualization, K.K.; supervision, L.P.; project administration, T.M.; funding acquisition, L.P. All authors have read and agreed to the published version of the manuscript.

## Funding

This research was funded by the National Research, Development and Innovation Fund of Hungary, grant number no. 2018-1.2.1-NKP-2018-00005 and by the Eötvös Lóránd Research network, grant number SA-82/2021.

## Informed Consent Statement

Not applicable.

## Data Availability Statement

The datasets supporting the conclusions of this article are included within the article.

## Conflicts of Interest

The authors declare no conflicts of interest.

### Abbreviations

The following abbreviations are used in this manuscript:

β-TrCP: Beta-transducin repeat-containing protein
C3H/10T1/2: Cell line isolated from C3H mouse embryo, exhibiting fibroblast morphology, functionally similar to mesenchymal stem cells
CHAPS: 3-[(3-cholamidopropyl)dimethyl-ammonio]-1-propane sulfonate
CK1: Casein kinase 1
DMEM: Dulbecco minimal essential medium
EC_50_: Half maximal effective concentration
EDTA: Ethylenediaminetetraacetic acid
GSK3β: Glycogen synthase kinase 3β
HEK: Human Embryonic Kidney
Hh: Hedgehog
NIH/3T3: Fibroblast cell line isolated from NIH/Swiss mouse embryo
PTCH1: Patched-1 protein
rhShh_1314-SH: Non-lipidated recombinant human sonic hedgehog protein expressed in *E. coli* rhShh_1845-SH Non-lipidated recombinant human sonic hedgehog protein expressed in *E. coli*; Cys24 residue has been replaced by a hydrophobic Met-Ile-Ile sequence to mimic the hydrophobic palmitoyl moiety of dually lipidated sonic hedge-hog protein
rhShh_8908-SH: Recombinant dually lipidated human sonic hedgehog protein expressed in HEK cells
rhWIF1: Recombinant human WIF1
Shh: Sonic hedgehog
WIF1: Wnt inhibitory factor 1

## Notes

### Competing Interest Statement

The authors have declared no competing interest.

